# EnsembleCNV: An ensemble machine learning algorithm to identify and genotype copy number variation using SNP array data

**DOI:** 10.1101/356667

**Authors:** Zhongyang Zhang, Haoxiang Cheng, Xiumei Hong, Antonio F. Di Narzo, Oscar Franzen, Shouneng Peng, Arno Ruusalepp, Jason C. Kovacic, Johan LM Bjorkegren, Xiaobin Wang, Ke Hao

## Abstract

The associations between diseases/traits and copy number variants (CNVs) have not been systematically investigated in genome-wide association studies (GWASs), primarily due to a lack of robust and accurate tools for CNV genotyping. Herein, we propose a novel ensemble learning framework, ensembleCNV, to detect and genotype CNVs using single nucleotide polymorphism (SNP) array data. EnsembleCNV a) identifies and eliminates batch effects at raw data level; b) assembles individual CNV calls into CNV regions (CNVRs) from multiple existing callers with complementary strengths by a heuristic algorithm; c) re-genotypes each CNVR with local likelihood model adjusted by global information across multiple CNVRs; d) refines CNVR boundaries by local correlation structure in copy number intensities; e) provides direct CNV genotyping accompanied with confidence score, directly accessible for downstream quality control and association analysis. Benchmarked on two large datasets, ensembleCNV outperformed competing methods and achieved a high call rate (93.3%) and reproducibility (98.6%), while concurrently achieving high sensitivity by capturing 85% of common CNVs documented in the 1000 Genomes Project. Given this CNV call rate and accuracy, which are comparable to SNP genotyping, we suggest ensembleCNV holds significant promise for performing genome-wide CNV association studies and investigating how CNVs predispose to human diseases.

## INTRODUCTION

As known GWAS loci only account for a fraction of disease/trait heritability (i.e., “Missing heritability”) (1), assessment of other types of human genetic variation besides single nucleotide polymorphisms (SNPs) is warranted. The human genome is rich in structural diversity, where copy number variants (CNV) are the most common form. CNVs are individually rare but collectively common across the human population (2). In fact, an estimated 8% of the general population carries a large (>500 kb) deletion or duplication occurring at an allele frequency of <0.05% (1,2). Moreover, CNVs affect transcription in mouse (3) and human (4), and contribute to a variety of different diseases (5). However, methods to genotype CNVs, particularly at the population level, are still in their infancy, lagging substantially behind genotyping of SNPs (6,7). Furthermore, the landscape of CNVs in the human genome is not fully characterized, including accurate assessments of CNV boundaries in terms of probes targeting each specific CNV on the SNP arrays. Taken together, although many CNV analysis software tools exist, the performance (sensitivity and accuracy) is sub-optimal, making CNV characterization significantly more challenging than SNP genotyping.

In general, existing CNV calling methods based on SNP array data can be categorized into two types: (i) individual-wise analysis and (ii) joint analysis of multiple individuals (see review by Pinto et al. (8)). Among individual-wise analysis, popular methods include hidden-Markov model (HMM)-based methods, such as PennCNV (9) and QuantiSNP (10), and segmentation-based methods, such as CBS (11) and fused-lasso methods (12,13). These methods utilize various types of information derived from SNP (or CNV) probes such as total intensity (i.e. Log R Ratio; LRR) and allele fraction (i.e., B Allele Frequency; BAF) from Illumina platforms as well as external information, such as population frequencies of the B allele at each locus (9) and linkage-disequilibrium (LD) structure between adjacent loci (14). Generally, these methods are good at detecting rare, large CNVs spanning at least ten probes on a SNP array. On the other hand, they rely on the assumption that allele intensities are properly normalized such that they are comparable across probes throughout the genome, and are thus less tolerant to spatial noise (e.g. genomic wave related to GC content) and heterogeneity across different loci (15). For example, at copy number polymorphism (CNP) loci with a high frequency of CNV alleles, the baseline LRR corresponding to a normal copy number may be distorted during normalization, and as a result, deviates from 0, making individual-based analysis error-prone. Moreover, from these methods, the individual-level CNV calls are frequently not aligned across individuals, adding additional difficulties in comparing CNVs in the downstream analysis.

Joint analysis of multiple individuals, such as iPattern (8), Piet (16), and msscan (17), to name a few, takes advantage of consensus CNV signals across individuals, which are particularly useful for CNP detection. They often align CNVs called across individuals into CNV regions (CNVRs), making the downstream analysis more accessible than individual-based methods. However, the boundaries of such constructed CNVRs are not carefully recalibrated. Moreover, they mainly focus on signal patterns in total intensity and do not fully utilize other CNV-related information (e.g., BAF), and as a consequence, are less sensitive to detect rare CNVs than individual-based methods. Lastly, both individual-wise and joint analyses usually report CNV calls only, and do not perform direct genotyping (i.e., explicitly differentiating between normal copy number and missing genotype). This fact, we believe, creates additional barriers for quality control and downstream association analyses.

In sum, many CNV calling methods have been proposed, each with various strengths and weaknesses. Thus, it is logical to aggregate multiple methods using an ensemble machine learning framework with the aim of achieving superior statistical performance. Herein, we propose a novel CNV detection and genotyping framework, ensembleCNV, which is primarily implemented in two phases: (1) the detection phase: initially locating CNVRs by assembling CNV calls from multiple methods with complementary advantages; (2) the re-genotyping phase: refining the initial calls with local models tuned for each CNVR. The ensembleCNV framework also includes steps to identify and eliminate batch effect at raw data level, which are essential to generate high-quality CNV signals. By leveraging large empirical datasets, we compare and intensively evaluated the performance of ensembleCNV with existing methods.

## MATERIAL AND METHODS

### Food Allergy (FA) dataset

In the genome-wide association study (GWAS) of food allergy (FA) in a US cohort of children with/without FA and their biological parents (18), a total of 2,790 blood DNA samples, including 839 nuclear families and 100 technically duplicated pairs, were genotyped on the Illumina HumanOmni1-Quad BeadChip with 1,048,713 SNP probes and 91,706 CNV probes. The mean and median of distances between neighboring probes are 2.63kb and 1kb (Supplementary Table S1). After quality control (QC), a total of 2,765 samples remained, including 835 nuclear families and 95 technically duplicated pairs (see Quality Control section for details). The majority (85.5%) of the samples are of European ancestry (Supplementary Table S1).

### STARNET dataset

In the Stockholm-Tartu Atherosclerosis Reverse Network Engineering Task study (STARNET) (19), a total of 874 blood DNA samples collected from coronary artery disease (CAD) patients, including 12 technically duplicated pairs, were genotyped on the Illumina HumanOmniExpressExome-8 BeadChip with 951,117 SNP probes. The mean and median of distance between neighboring markers are 3.23kb and 1kb (Supplementary Table S1). After quality control (see Quality Control section for details), a total of 834 samples remained, including 12 technically duplicated pairs (Supplementary Table S1).

### 1000 Genomes Project (KGP) CNV dataset

We downloaded the KGP structural variant (SV) dataset (in GRCh37 coordinates) in variant call format (VCF), which consists of 68,818 SVs detected by whole-genome sequencing (WGS) in 2,504 individuals from different genetic populations (20). In this study, we focused on the subset (CNV dataset) of 40,975 bi-allelic deletions (DELs), 6,025 bi-allelic duplications (DUPs) and 2,929 multi-allelic copy number variants (CNVs) (Supplementary Table S2), because only these types of SVs can be detected by SNP array platforms. To compare with the CNVs detected in the FA and STARNET SNP array data, we further defined the subset of detectable CNVs spanning at least 5 probes of the SNP arrays used in the two studies, resulting in a total of 6,456 and 3,571 detectable CNVs for the FA and STARNET datasets, respectively (Supplementary Table S2).

### Data processing and quality control

We processed the raw data (i.e., .idat files) with Genome Studio 2011.1 (Illumina, CA) following similar protocols as described (21). For data processing with Genome Studio, an important step is to update the cluster centers corresponding to the so-called AA, AB and BB genotypes for each SNP probe (CNV probe has only one cluster), where A and B refers to the two alleles for the SNP probe. This was done by re-clustering on the data points from samples with high call rate (e.g., >95%). This step is necessary since the cluster centers (in .egt file) accompanying the SNP array, or which are built from other studies, do not always align with the real cluster centers in the current study. In the FA and STARNET datasets, the sample sizes are large enough to build customized clusters. For the updated cluster centers of each probe, we updated genotype calls and derived quantities (including LRR and BAF used for CNV analysis) and exported them to final reports.

We performed two types of sample-level quality control (QC). Firstly, as routine QC for a GWAS based on SNP genotype data, we excluded samples with (1) mismatched genders, (2) excessive missing genotype rates, (3) excessive heterozygosity in autosomes (an indication of potential sample contamination) and (4) outliers in principle component analysis (PCA). We retained technical duplicates in this study to evaluate CNV methods (see below). It should be noted that we did not perform QC at the probe-level (as in a typical QC for GWAS), because SNP probes with high missing genotype rates and deviation from Hardy-Weinberg equilibrium may be associated with CNVs. At these SNP probes, data points corresponding to CNVs usually deviate from empirical genotype clusters and such deviation is especially magnificent for those with CN = 0 (*i.e.*, 0 copies of the allele) (13,14). Secondly, we performed additional sample-level QC to remove samples with abnormal CNV signals and alleviate batch effects. Please refer to ensembleCNV step (b) (see below) and Supplementary Results for details.

### ensembleCNV workflow

The overall workflow of ensembleCNV is summarized in Figure 1. This involves an initial localization of CNVRs by assembling CNV calls from multiple methods with complementary advantages, and then a refining of the initial calls using local models tuned for each CNVR. The workflow consists of two major phases implemented in five steps. In the initial detection phase, we processed the raw image data with Genome Studio (Illumina, CA) and extracted genotype and CNV signals, particularly log R ratio (LRR) and B allele frequency (BAF) from the final report. Next, we applied three popular CNV detection methods, PennCNV (9), QuantiSNP (10), and iPattern (8), to create the initial CNV call sets, respectively (step (a)). In the raw data and initial call sets, batch effects may exist and affect the downstream analysis. For this reason, we applied PCA on the raw LRR data and sample-level summary statistics of CNV results from the individual callers to identify batches. When batches existed, we re-processed each batch along the pipeline for the initial detection phase (step (b)). In the following ensemble and re-genotyping phase, we aimed to address three sub-tasks: First, we aligned the CNVs called from the individual methods in all subjects and merged overlapping CNVs to construct initial CNVRs using a heuristic algorithm (step (c)). Second, we trained a local likelihood model on both LRR and BAF CNV signals at each CNVR with the “global” information from frequent CNVRs incorporated (see Supplementary Methods); and assigned a copy number (CN) genotype to each subject, accompanied by a genotyping quality (GQ) score to quantify the confidence level (step (d)). Third, we refined the boundaries of CNVRs using the correlation structure of the LRR values of the probes around the CNVRs (step (e)). The whole pipeline results in CN genotype data of refined CNVRs across each subject. Details of the implementations are described as follows.

**Figure 1:**
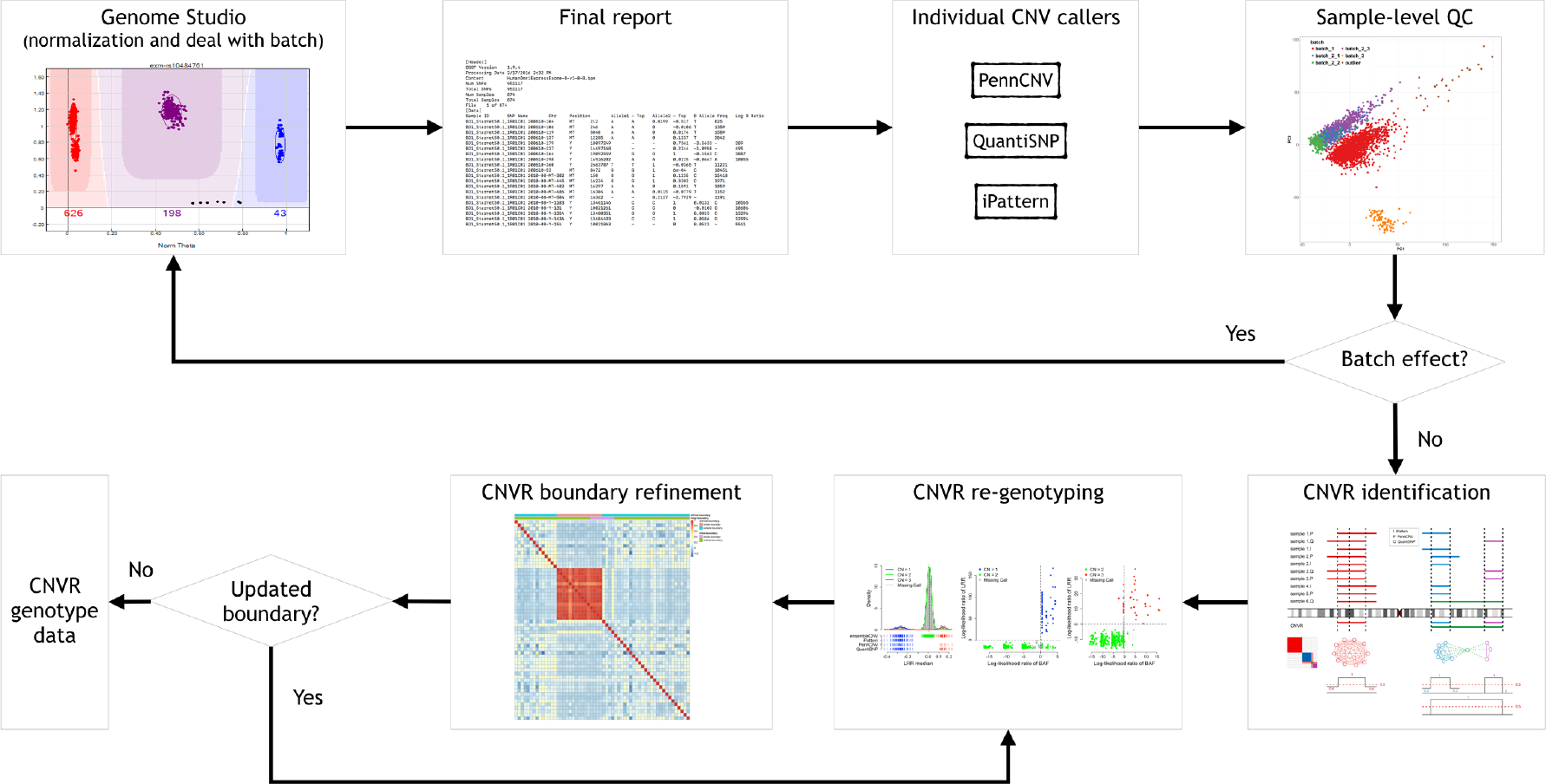
Workflow of ensembleCNV.

#### (a) Initial CNV calls by selected methods

We chose PennCNV (9), QuantiSNP (10), and iPattern (8) to make the initial CNV calls using final report files generated by Genome Studio 2011.1 (Illumina, CA). These methods use different information to call CNVs from different perspectives. Both PennCNV and QuantiSNP are hidden Markov Model (HMM)-based approaches, which take LRR (reflecting total copy number) and BAF (reflecting allelic proportion/balance) jointly as observed data while modeling copy number (CN) status in the hidden layer. PennCNV further accounts for population frequency of the B allele, which can be estimated by the data. They make CNV calls on an individual-wise basis. iPattern takes normalized intensities from fluorescent measurements of the two alleles of each SNP as input, calculates total intensities for each SNP, and normalizes the total intensities (reflecting total CN) across individuals. It then screens the genome with sliding windows, within which a Gaussian mixture model (GMM) is fitted to the normalized total intensities across individuals and CNV calls are made based on the fitted model. While PennCNV and QuantiSNP are good at calling large rare CNVs in a more sensitive way, iPattern performs better in calling more frequent CNVs across individuals. We took the complementary advantages of these methods and combined their discovery sets to boost the sensitivity of CNV detection for the initial call set.

#### (b) Sample-level QC for batch effects

We used two orthogonal signals to identify batch effects in CNV calling: (i) Along with CNV calls, the three detection methods can generate per-sample summary statistics, such as standard deviations (SD) of LRR, SD of BAF, wave factor in LRR (15), BAF drift (9), and the number of CNVs detected, reflecting the quality of CNV calls at the sample level. Since these quantities are highly correlated among themselves and between methods (Supplementary Figure S1), we used PCA to summarize their information. By examining the first two or three PCs, we can identify sample outliers or batches that deviate from the majority of the normally behaved samples (Supplementary Figure S2). (ii) Batch effects may also be orthogonally reflected in the first two or three PCs from the LRR data before any CNV analysis is performed. We randomly selected 100,000 probes and applied PCA to the LRR values at these probes across individuals and visualized the first few PCs in scatter plots. Batches can be identified by visual check if they exist (Supplementary Figure S2). The batches identified by these two independent approaches should be consistent with each other. While isolated outliers were excluded from downstream analysis, if batch effects were identified, we re-normalized the samples within each outstanding batch with Genome Studio (see data processing section), re-did the CNV calling in step (a), and combined the re-called CNVs with the remaining call set of good quality (Figure 1). Please refer to Supplementary Results and Figures S1-3 for details regarding the identification and removal of batch effects in the FA dataset.

#### (c) Construction of CNVRs

We defined CNVR as the region in which CNVs called from different individuals by different callers substantially overlap with each other. CNV events belonging to the same CNVR are comparable across individuals and thereby the estimation of population frequency can be made. For copy number polymorphisms (CNPs) frequently observed in populations and inherited CNVs segregating within pedigrees, their boundaries would be aligned precisely across individuals. For recurrent CNVs (but not CNPs), their affected genomic regions would largely overlap, albeit if not exactly aligned.

We modeled the CNVR construction problem as identification of cliques (a sub-network in which every pair of nodes is connected) in a network context (Supplementary Figure S4), where (i) CNVs detected for each individual from a method are considered as nodes; (ii) two nodes are connected when the reciprocal overlap between their corresponding CNV segments is greater than a pre-specified threshold (e.g. 30%); (iii) a clique corresponds to a CNVR in the sense that, for each CNV (node) belonging to the CNVR (clique), its average overlap with all the other CNVs of this CNVR is above a pre-specified threshold (e.g. 30%). The computational complexity for clique identification can be dramatically reduced in this special case, since the CNVs can be sorted by their genomic locations and the whole network can be partitioned by chromosome arms – CNVs from different arms never belong to the same CNVR. Correspondingly, the adjacent matrix representing the network model is banded along the diagonal (Supplementary Figure S4).

Briefly, for CNVs in each chromosome arm, which were sorted by their genomic locations, we constructed CNVRs in a forward-screening and backward-pruning procedure. In the forward-screening step, we initialized the first CNVR with the first CNV in the list and screened the remaining CNVs in their genomic order. For the current CNV under consideration, we compared it against all existing CNVRs. If its average overlap with all CNVs of a most overlapping CNVR was above the pre-specified threshold (see above definition (iii)), we assigned it to this CNVR; otherwise we created a new CNVR with it. The screening continued until all CNVs in the list were assigned to a CNVR. In the backward-pruning step, for each CNVR, we re-checked, for each CNV belonging to this CNVR, if its average overlap with all the other CNVs of this CNVR was above the pre-specified threshold. If a CNV did not meet this criterion, we removed it from this CNVR. This pruning procedure would continue until no more CNVs could be removed from the CNVR. The leftover CNVs would be assigned to other CNVRs whenever possible; otherwise, they would be assigned to newly created CNVRs.

To define the boundary of each CNVR, we stacked all CNVs in a CNVR and defined the footprint carried by at least a pre-specified proportion of CNVs (e.g. 50%) as the initial boundaries of this CNVR (Supplementary Figure S4). We adopted this major-vote type of strategy to enrich CNV signals within the CNVR and reduce the noise from surrounding probes. The initial boundaries were refined in an iterative way as shown in Figure 1 and step (e) below.

#### (d) CNV re-genotyping

The initial CNV calls within a CNVR may be mixed with false positives and false negatives from the initial call set. Moreover, the baseline LRR value corresponding to normal CN status may substantially deviate from 0, violating the essential model assumptions for individual-wise CNV callers (e.g., PennCNV and QuantiSNP) (Supplementary Figures S5A and S6). To address these issues, we re-genotyped CN status per individual at each CNVR by a locally fitted likelihood model, with information from other CNVRs borrowed for the initialization of model parameters (see Supplementary Methods). Both the LRR and BAF signals from SNP probes and the LRR signal from CNV probes within a particular CNVR were used for model fitting. With the additional signal from BAF, the original samples with non-identifiable CN status based on the LRR signal alone (e.g., iPattern) may be distinguishable in the expanded BAF-LRR 2-D space (Supplementary Figure S5B-C).

For BAF signal, denote *X*_*ij*_ (*i* = 1,…, *n*; *j* = 1,…, *p*) as the observed BAF value for the *i*-th individual at the *j* -th SNP probe in the CNVR. We adopted the mixture model (9):

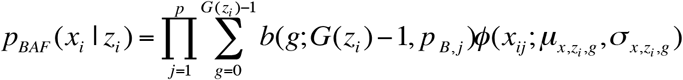

where *b*(*g*; *G*(*z*) −1, *p*_*B*_) and *ϕ*(*x*; *µ*_*x,z,g*_,*σ*_*x,z,g*_) are density functions for binomial and normal distributions respectively; *z*_*i*_ (*z*_*i*_ ∈ {0,1,2,3}) is CN; *G*(*z*_*i*_) is the number of possible genotypes associated with the CN (e.g., *G*(*z*_*i*_ = 3) = 4 corresponding to genotype AAA, AAB, ABB and BBB); *p*_*B*_ is the population B allele frequency (PFB), which can be retrieved from PennCNV analysis (9) in step (a). The parameters in the BAF model are estimated from a set of selected CNVRs (see Supplementary Methods).

For LRR signal, denote the median value of all probes within the CNVR as *y*_*i*_ (*i* = 1,…, *n*) for the *i* -th individual. We adopted the commonly used Gaussian mixture model (GMM):

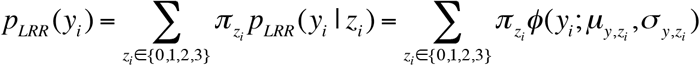

The model can be fitted by an EM algorithm (22). For a CN Gaussian component in the GMM with adequate sample size (e.g. >= 10) from the initial call set generated in step (a), we use these samples to estimate the initial value of parameters for the EM algorithm; otherwise, we adopted the parameters estimated from a set of selected CNVRs as the initial value (see Supplementary Methods). It should be noted that LRR signals are not always properly normalized and centered at so we used the mode of *y*_*i*_ of the samples absent from the initial CNV call set (i.e., CN = 2) for the CNVR to estimate the initial location of normal CN component (*µ*_*y,z*=2_) and calculate the relative locations of other CN components with respect to *µ*_*y,z*=2_.

Once the CNVR-specific model is fitted, we computed the Phred-scaled likelihood (PL) of a CN status (*z*_*i*_ ∈ {0,1,2,3}):

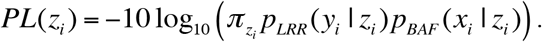

We decided the CN genotype (including CNV and normal CN) as the one with the smallest PL value, with associated genotype quality (GQ) score defined by the difference between the smallest and the second smallest PL values. This definition is similar to the GQ score used in GATK pipeline for the analysis of next-generation sequencing data (23). The GQ score can be used to measure the confidence of a CN genotype call. The CN genotype with GQ score greater than a pre-specified threshold will be reported and otherwise be set as missing (“no call”). The GQ threshold can be tuned in a data-driven manner that balances accuracy and call rate. For example, technical duplicates are often adopted for QC purposes. We can adjust the GQ score threshold to a level that the CNVs detected in technical duplicates achieve a high concordance rate while the call rates at the individual-level and CNVR-level are not heavily compromised (see Results).

#### (e) Boundary refinement

For a CNVR with high frequency of variant alleles, the LRR signals are highly correlated across individuals among involved probes (Supplementary Figure S7). We can take advantage of this structure to further refine CNVR boundaries. In other words, we are able to find a sub-block of high correlations within a local correlation matrix. This strategy has also been adopted for CNV detection in other studies (24,25). Considering the ***M*** SNPs within the local region by expanding the initial boundaries several times (e.g. twice the size of the initial region on both sides), let *r*_*ij*_ be the Pearson correlation of LRR values between SNP *i* and *j*. The refined left *l̂* and right *r̂* boundaries are obtained by optimizing:

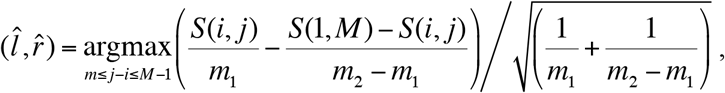

where 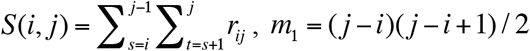, *m*_2_ = *M*(*M* – 1)/2, and *m* is the pre-specified minimum size of CNVR (e.g., spanning 5 probes). This optimization is feasible by simple exhaustive search, since the number of SNPs ***M*** involved in a CNVR is commonly within the range of tens to hundreds.

When the refined boundaries are different from the initial boundaries, the probes falling within the range of updated boundaries will change. We needed to update the local likelihood model as in step (d) and re-did the genotyping step (Figure 1). If several CNVRs share the exact boundaries after boundary refinement, we merged them into one.

### Concordance rate in duplicated pairs

To avoid overestimating the concordance rate, for each pair of technical duplicates, we only considered the CNVRs with CNV genotype (CN ≠ 2) in at least one of the pair, while those with only normal CN genotype (CN = 2) in both were excluded. The concordance rate was defined as the proportion of CNVRs with consistent CNV status in both duplicates among all the CNVRs considered for each pair of duplicates.

### Mendelian errors and transmission rate in trios

From the nuclear families in the FA cohort, we included those with complete trios (father-mother-child). Larger nuclear families with multiple children were converted to multiple trios. For example, a quartet family was converted to two trios, each with one child and the parents. This resulted in a total number of 1019 trios. In each trio, the CNVRs with normal CN status (CN = 2) in all three individuals were excluded. A CNVR has a Mendelian error if the CN status in the trios does not follow the normal inheritance pattern. For example, if CN = 0 in the father, CN = 2 in the mother, then CN = 2 in the child will lead to a Mendelian error. A Mendelian error could possibly arise from a *de novo* mutation in the child, false positive CNV calls in the child, or false negatives in the parents. To avoid ambiguity in the estimation of transmission rate in a trio, we only considered the CNVR with CNV status (CN ≠ 2) in only one of the parents. The transmission rate was estimated as the proportion of CNVRs carrying CNV genotype in the child among all CNVRs considered in the trio.

### Sensitivity analysis

We used the “detectable” subset of the 1000 Genomes Project (KGP) CNV dataset (Supplementary Table S2) as the reference to evaluate the sensitivity of the six methods in CNV detection. A CNV in KGP data was considered as detected by a method if a CNV from the call set of the method was found with ≥ 30% reciprocal overlap with the KGP CNV and the allelic types of the two CNVs were compatible. For example, if the CNV called by the method had only a deletion allele while the KGP CNV had only a duplication allele, then the KGP CNV was not counted as detected even though they had ≥ 30% reciprocal overlap. The sensitivity of a method was defined as the proportion of the CNVs in the KGP data that could be detected by the method.

## RESULTS

### Performance of ensembleCNV

We evaluated the performance of ensembleCNV using two empirical datasets from the FA and STARNET studies (see Material and Methods; Supplementary Table S1) and compared with the three methods we adopted in our pipeline. In addition, we also considered two simple integration methods commonly used in CNV studies: (i) the “intersection” method where the CNVs called by at least two of the three methods (i.e., major voting) were selected to the final call set; and (ii) the “union” method where the CNVs called by any of the three methods were added to the final call set (Supplementary Figure S8). For the other five methods, we used the ensembleCNV CNVR construction algorithm to create CNVRs. Key summary statistics of the CNV call sets from the six methods are shown in Supplementary Table S3. The individual performance of these six methods were compared from three perspectives: (1) concordance rate of detected CNVs between technical duplicates in both datasets; (2) Mendelian error and transmission rate evaluated in trios of FA study; (3) the quality of detected CNVs evaluated by the external CNV data from the 1000 Genomes Project (KGP) (20).

### Concordance rate and genotype call rate

To benchmark the accuracy of CNV detection methods, the ideal way is to use real datasets with known ground truth of CNVs in all subjects. Such a benchmark is usually not available, especially for large-scale genetic studies. However, for QC purposes in SNP genotyping, technical duplicates are often considered in the experimental design as in the FA and STARNET studies. Thus, the concordance rate (see Methods for definition) of CNV calls in duplicated pairs, i.e. the reproducibility, can be used as a surrogate of accuracy measurement – with high reproducibility being an indication of good quality for a CNV call set. In ensembleCNV, we also defined the genotyping quality (GQ) score (see Methods) to quantify the confidence of the CN genotype assigned to each individual at each CNVR. As the GQ score threshold increases, the concordance rate constantly increases with the median value across duplicated pairs gradually approaching 100% at the cost of decreased sample-wise and CNVR-wise call rates (Supplementary Figures S9 and S10). To achieve a balance between concordance rate and call rate, we selected GQ score thresholds of 15 and 20 in the FA and STARNET datasets, respectively (Supplementary Figures S9 and S10). We also used these thresholds in the results sections below. Of note, the strategy of utilizing technical duplicates along with the GQ score can be used when applying ensembleCNV in real data analyses. For the other five methods, however, there is no such quantification of genotyping confidence for every individual at each CNVR. In particular, these methods do not distinguish between normal CN (CN = 2) vs. no call. Instead, we set the genotype of an individual without a CNV call (CN ≠ 2) at a CNVR as normal (CN = 2). The call rate, therefore, cannot be defined for these five methods.

In this evaluation, not only did ensembleCNV achieve the highest concordance rates with medians of 98.6% and 95.5% in the FA and STARNET datasets, but it also resulted in the greatest stability (i.e., the smallest variability) across duplicated pairs (Figure 2A and D). On the other hand, the medians of sample-wise call rate and CNVR-wise call rate reached 93.3% and 97.0% for the FA data and 96.3% and 99.5% for the STARNET data, respectively (Figure 2B, C, E, and F). A similar QC based on call rate at both individual and CNVR levels can be performed as what is routinely done for SNP QC in GWAS. For the purpose of fair comparison, we kept all samples and CNVRs generated by ensembleCNV without any filtering unless otherwise specified. Following ensembleCNV, the joint analysis based method, iPattern, performed better than the individual-wise analysis based methods, PennCNV and QuantiSNP, in terms of concordance rate. It should be noted that the straightforward integration methods “intersection” and “union” did not make any improvement on the individual methods, but instead reached a compromise between the three methods – they underperformed as compared to iPattern and outperformed as compared to PennCNV and QuantiSNP (Figure 2A and D). Moreover, in the FA data, the concordance rate produced by ensembleCNV was mostly comparable between the duplicated pairs, either within the same batches or those belonging to different batches. This suggests robustness of ensembleCNV to batch effects, whereas the other five methods were vulnerable (Supplementary Figure S11).

**Figure 2:**
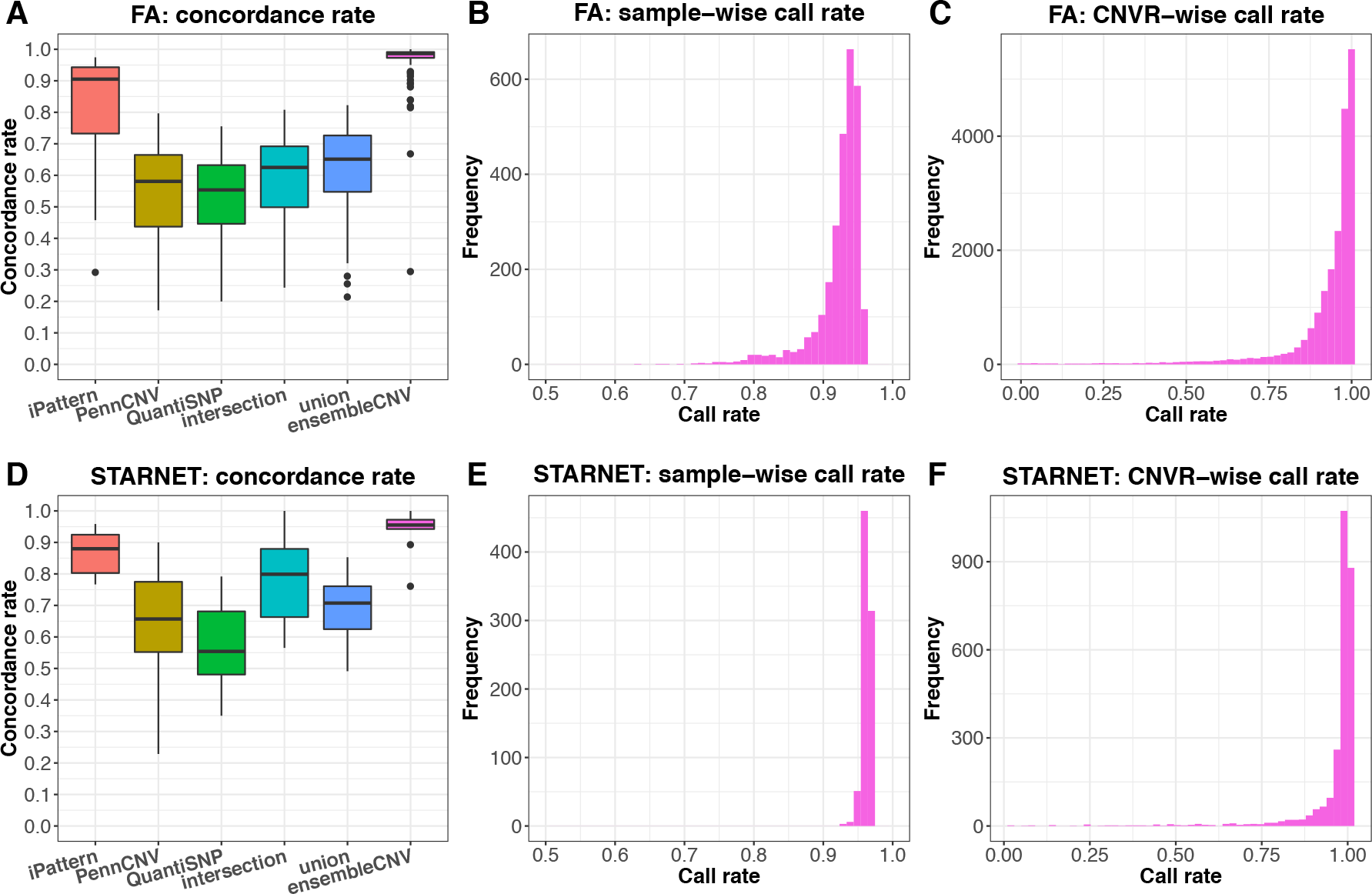
Concordance rate and call rate of detected CNVs. Results are summarized for the FA (A-C) and STARNET datasets (D-F), respectively. In (A) and (D), the concordance rate of CNV genotype for technically duplicated pairs are compared among six methods. In (B), (C), (E) and (F), the distribution of the sample-wise and CNVR-wise call rates of CNV genotype are shown for ensembleCNV, since only ensembleCNV makes direct genotyping in each CNVR and distinguishes between normal copy number and missing genotype.

### Mendelian error and transmission rate

Taking advantage of family information from the FA dataset, we calculated the number of Mendelian errors in the trios (see Methods). Though a Mendelian error may imply a *de novo* CNV, the number of such errors is expected to be within a certain range. As suggested by KGP CNV data (Supplementary Table S2), the median number of singleton CNVs per sample is 8. Therefore, an excessive number of Mendelian errors per trio indicates poor quality of a CNV call set. Figure 3A summarizes the number of Mendelian errors per trio for the six methods. The median number for ensembleCNV is 13, a little above the number suggested by KGP data. It should be noted that the GQ score threshold was not optimized for Mendelian errors (see above section). With the increment of GQ score threshold, the number of Mendelian errors continues to decrease at the expense of call rate (Supplementary Figure S12). In contrast, the median number for iPattern is 45 and those numbers for the other four methods are all above 200, implying a large amount of false positive CNV calls in offspring and/or false negatives in parents. Concurrently, with trio data, we estimated the transmission rate of CNVs from parents to offspring (see Methods). Normally, the average transmission rate should be around 0.5 (26). The medians of transmission rates in the trios for ensembleCNV and iPattern were close to 0.5, whereas the median values for the other four methods were all well below 0.5 (Figure 3B). Moreover, as expected, the median of transmission rates for ensembleCNV converged to 0.5 as the GQ score threshold increased (Supplementary Figure S12).

**Figure 3:**
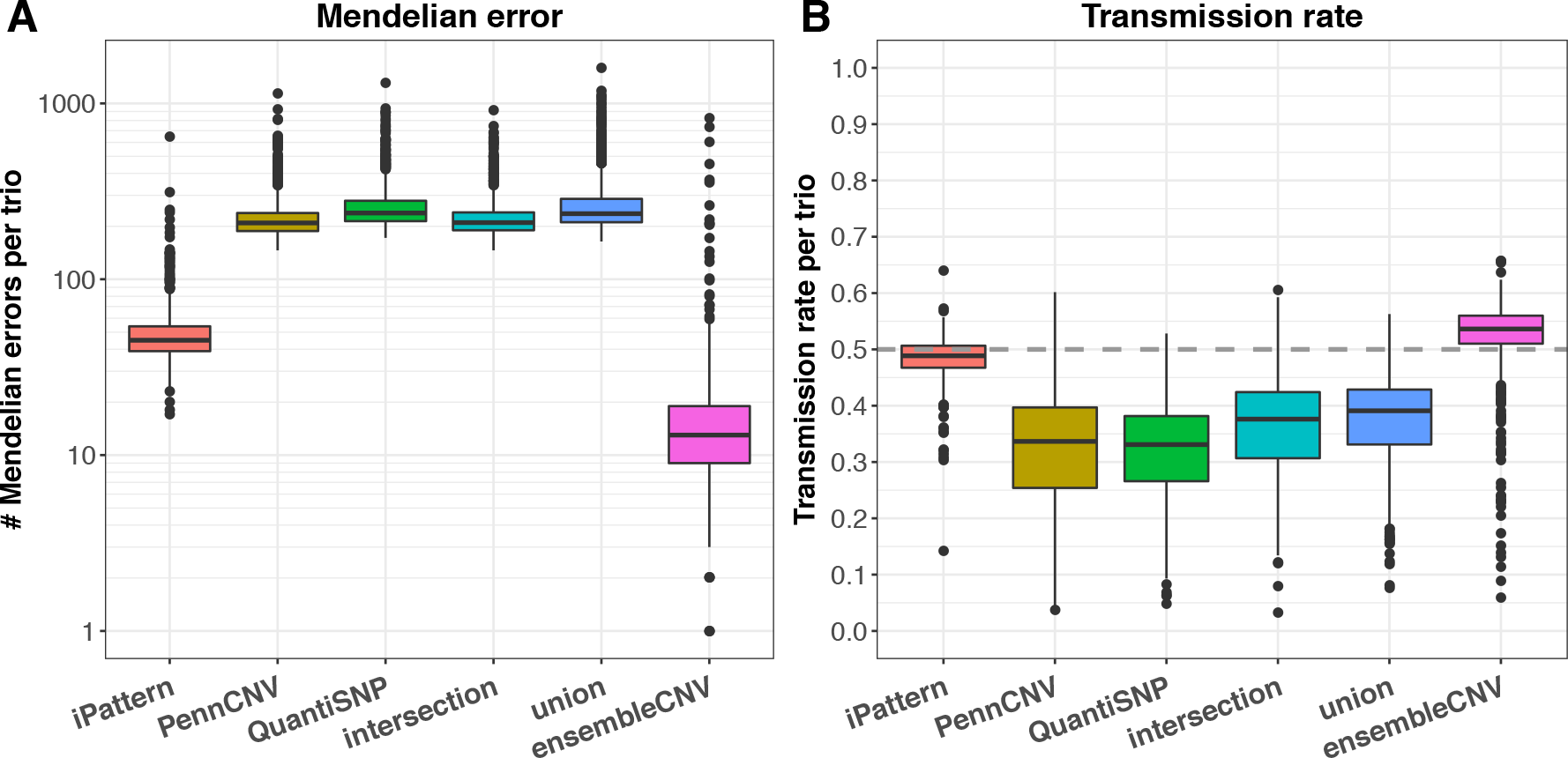
Mendelian error and transmission rate in FA trios. Given the information regarding nuclear families in the FA dataset, the Mendelian error (A) and transmission rate (B) calculated from CNV genotype data in trios were compared among the six methods. The y-axis in (A) is on log10 scale.

### Evaluation with 1000 Genomes Project CNV (KGP) data

We used the KGP CNV data (20) as an external dataset to evaluate the sensitivity of the six methods (Supplementary Table S2). Since KGP CNV data was produced by whole-genome sequencing (WGS), the size of detectable CNVs can be smaller than 1kb at base-pair resolution, which is beyond the capability of SNP array platforms. To make the reference data comparable to the results from the FA and STARNET datasets, we defined the subsets of “detectable” CNVs as those spanning at least 5 probes in the SNP array used in the FA and STARNET studies, respectively (Supplementary Table S2). Since the majority of subjects in the FA and STARNET studies are of European ancestry and the SNP arrays used (Supplementary Table S1) are mainly designed for European populations (reflected in the number of detectable singleton CNVs per sample; Supplementary Table S2), we focused on the statistics in European populations from KGP. The sensitivity assessment was further stratified by the allele frequency in KGP European populations at 1% (Figure 4). Overall, the total number of CNVs detected per sample from ensembleCNV (median: 620 and 38) was the closest, among the six methods, to the number (median: 633 and 48 for European populations) of the detectable subsets of KGP data in both the FA and STARNET studies (Supplementary Tables S2 and S3). Regarding the sensitivity, ensembleCNV was able to identify 85% and 71% of detectable common CNVs in the FA and STARNET data, respectively. That the sensitivity for rare CNVs was found to be much lower is not surprising (Figure 4). The call set from iPattern is the most conservative and biased toward common CNVs, while the other four methods tend to be slightly more sensitive than ensembleCNV at a greater cost of accuracy (Figures 2-4). Interestingly, for the detectable common CNVs (containing ≥5 probes on the SNP genotype array), the CN genotype frequencies documented in the KGP CNV dataset were better matched with those estimated by ensembleCNV than the other 5 evaluated methods (see Methods; Supplementary Figure S13).

**Figure 4:**
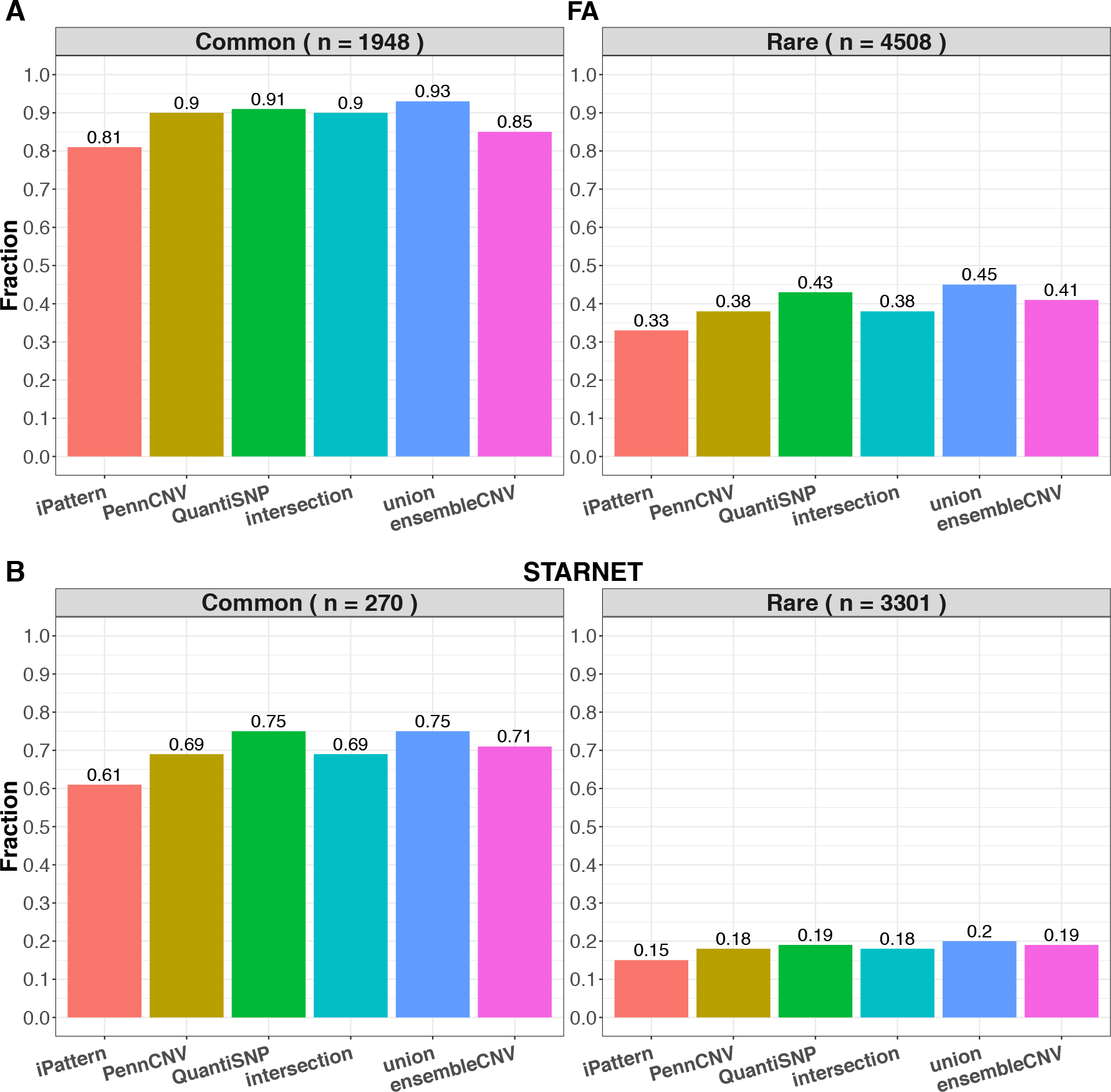
Sensitivity of detecting CNVRs in the KGP dataset. The fraction of detectable KGP CNVRs spanning at least 5 probes in the FA (A) and STARNET datasets (B) were compared among the six methods. In each subfigure, the detectable CNVRs are further stratified by allele frequencies in European populations of KGP into common (frequency > 1%; left panel) and rare CNVRs (frequency < 1%; right panel).

### Size and frequency distribution of CNVRs

The size and frequency of CNVs called by the different methods were found to be comparable both to each other and to CNVs released by KGP (Figure 5). In the FA study, 84.2% and 91.1% of all CNVRs were below the frequencies of 1% and 5%, respectively. In the STARNET study, 83.7% and 93.7% of all CNVRs were below frequencies of 1% and 5%, respectively. This frequency spectrum is similar to that observed for SNPs (27). Importantly, in the FA (by ensembleCNV) and KGP data, a total of 1,752 and 1,948 CNVRs with frequency ≥ 1% was detectable, offering sufficient statistical power to perform large CNV-GWAS.

**Figure 5:**
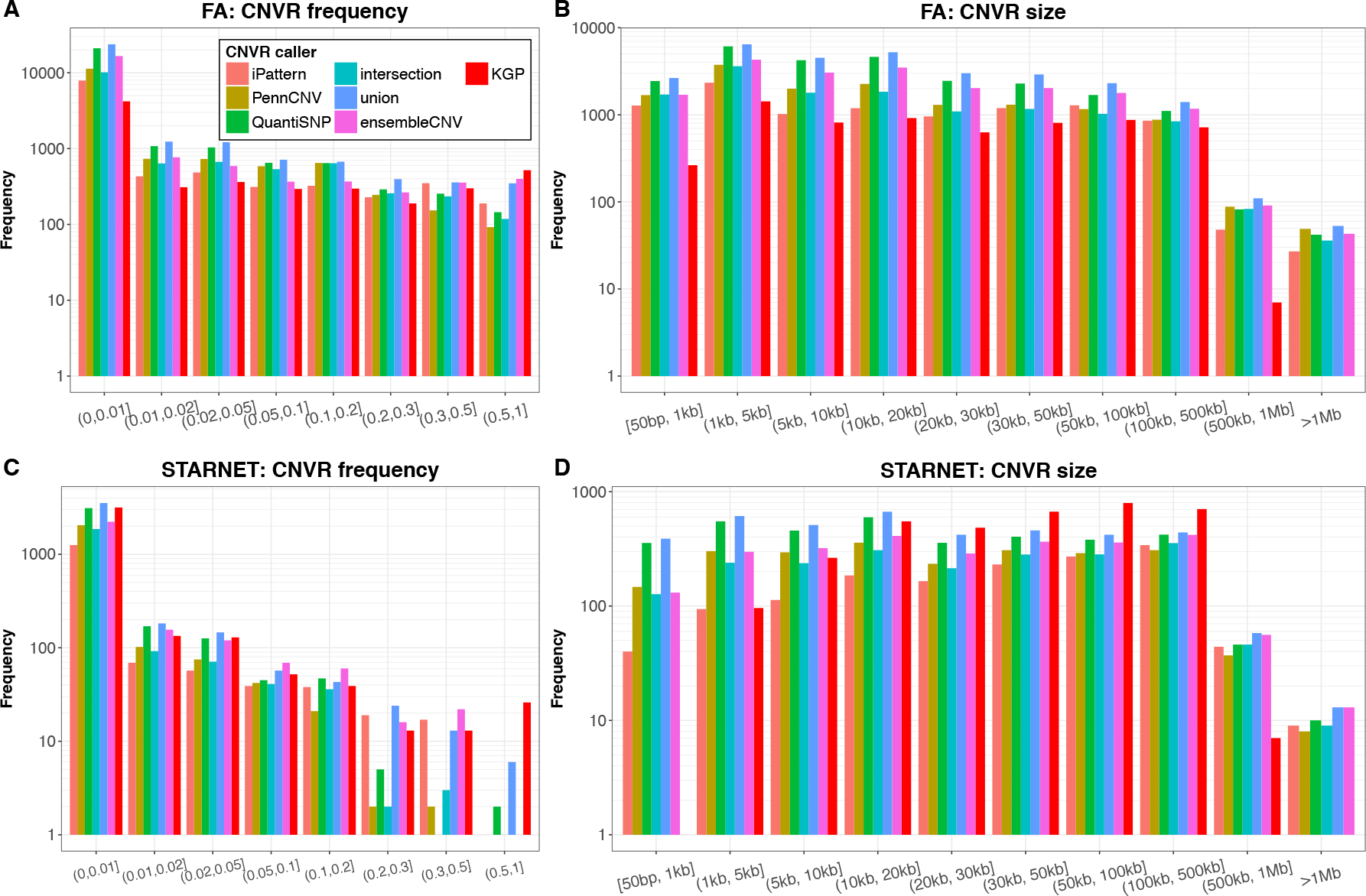
CNVR frequency and size distribution. The frequency (A and C) and size (B and D) of CNVRs detected in the FA (A and B) and STARNET (C and D) datasets using the six methods as well as those detectable CNVRs (spanning at least 5 probes in each of the dataset). The frequency of a CNVR refers to the proportion of the unrelated carriers of a CNV genotype instead of allele frequency at the CNVR. The frequency in KGP was calculated based on European populations.

The CNVRs detected had a wide size range (Figure 5B and 5D), which is consistent with findings in KGP. 63.7% and 43.8% of CNVRs in the FA and STARNET data were 20kb or shorter, respectively. It should be noted that there were only 2 CNVs with a size of greater than 1Mb in the KGP CNV dataset (Supplementary Table S2), and these two CNVs did not pass the 5-probe lower limit and thus did not appear in the detectable sets for the FA and STARNET studies.

### Correlation of CNVRs and nearby SNPs

A key question for CNV-GWAS is whether the CNVs are already well tagged by nearby SNPs, in which case making CNV calling becomes redundant. We estimated linkage disequilibrium (LD; r^2^) between each CNVR (with >90% call rate from ensembleCNV) and SNPs within 500 kb (Figure 6). In the FA data for all CNVRs, only 7.3% were tagged at r^2^ ≥ 0.6. In contrast, for CNVRs at frequencies ≥ 1% and ≥ 5%, the corresponding numbers were 46.5% and 76.3%, respectively. In STARNET, a SNP array with fewer SNPs than FA was used, and the LD between CNVRs and SNPs was therefore even lower. For all CNVRs, 4.4% were tagged at r^2^ ≥ 0.6, and for CNVRs at frequencies ≥1% and ≥5%, 15.3% and 36.9% were tagged at r^2^ ≥ 0.6, respectively. These results indicate that testing CNV-disease associations is necessary and potentially fruitful for both rare and common CNVs.

**Figure 6:**
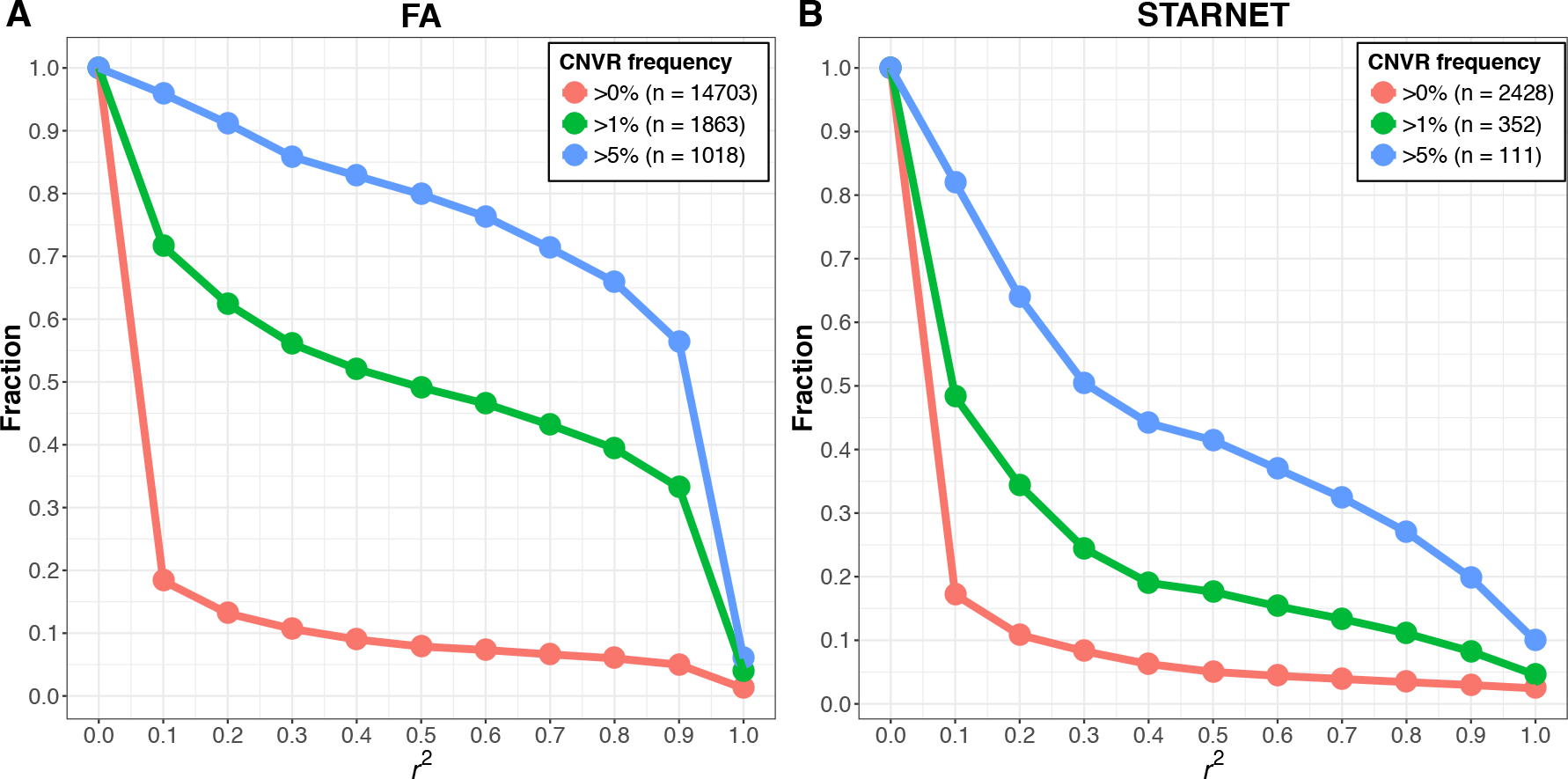
Linkage disequilibrium (LD) between CNVR and nearby SNP probes. LD (r^2^) was estimated as the maximum of squared Pearson correlations between the copy number at each CNVR resulted from ensembleCNV and the genotype (coded as 0, 1, and 2 copies of alternative alleles) of SNPs within 500kb upstream and downstream of the CNVR in the FA (A) and STARNET datasets (B). The plots show the fraction of CNVRs as the threshold of LD measure increases. The call set of CNVRs is further categorized by frequency greater than 0%, 1% and 5%, respectively, with the total number of CNVRs in each subset indicated in the legend. The frequency of a CNVR refers to the percentage of the unrelated carriers of a CNV genotype at the CNVR.

### Functional relevance of CNV

In the STARNET and FA datasets, we found that 5.58% and 17.49% of the genome were affected by CNVs, respectively. For more frequent CNVs (e.g. frequency ≥ 5%), 0.74% (FA) and 1.23% (STARNET) of the genome were affected. Furthermore, 0.54% (FA) and 0.4% (STARNET) of the genome were affected by CNVs with null (CN = 0) genotype.

Given that CNVs directly change the dosage of genes, they are likely functional and possibly important for disease. For that reason, we overlapped the CNVRs with the NHGRI-EBI GWAS catalog (28), holding 43,927 unique variants with convincing evidence of variant-trait associations. Importantly, 23.97% (10,530) of the GWAS catalog variants were affected by CNVRs (i.e., within CNVR boundaries), where 2% (881) were affected by frequent CNVRs and 1.1% (485) by CNVRs with null genotype.

It is known that most genomic loci can affect several traits either by affecting multiple genes at the locus or genetic pleiotropy (29-31). In our results, risk loci of the GWAS catalog affected by CNVs we detected in either the FA or the STARNET data were linked to multiple traits. The top diseases/traits with the most GWAS SNPs affected by CNVs were breast cancer (220 CNV-affected variants), schizophrenia (220 variants), obesity-related traits (213 variants), height (174 variants), and body mass index (170 variants). As an example, the 7q36.3 locus with susceptibility to testicular germ cell tumor (leading SNP rs11769858; p-value = 2e-8) (32) was within a CNVR (CNVR_954_r1_chr7_q) of 4kb length (Supplementary Figure S14A). In the FA cohort, 2014, 483 and 31 subjects are of 2, 1 and 0 copy numbers, respectively, at this CNVR (Supplementary Figure S14B). This CNVR (CNVR_954_r1_chr7_q) is reliably called based on 15 probes on the Illumina HumanOmni1-Quad-v1 BeadChip (Supplementary Figure S14C and D). The promoter region of a gene proposed to be responsible for testicular germ cell tumor risk (32), NCAPG2, was affected by this CNVR (Supplementary Figure S14A).

## DISCUSSION

Despite the availability of large, relevant datasets (e.g. SNP array of large GWASs) (7), the hypothesis that CNV broadly influences disease risk in the population and across diseases has not been systematically evaluated in a well-powered study. The main challenge has been a lack of robust and accurate methods to quantify CNVs. Here we describe and report on the utility of ensembleCNV for detecting and genotyping CNVs; a technique that is readily applicable to a large amount of existing data. We demonstrate that our ensemble approach quantifies CNV genotype with accuracy and properties comparable to SNP genotyping, and as such paves the way for large-scale population-based CNV-disease association studies (i.e., CNV-GWAS).

EnsembleCNV has several key advantages over existing methods:

1. *High genotyping accuracy and reproducibility*. In the STARNET and FA cohorts, we observed 96.2% and 98.6% consistency, respectively, among technical duplicates (Figure 2), and the median number of Mendelian errors was 13 per family in the FA data, which outperformed other methods.
2. *Detection and re-genotyping functions*. Many existing methods are designed as CNV detection tools (e.g. PennCNV and QuantiSNP) and do not distinguish CN=2 vs. “no call”, causing substantial mis-classification. EnsembleCNV has both detection and re-genotyping functions and employed a GQ score to identify “no call” for genotypes with low confidence.
3. *High genotyping call rate*. In the STARNET and FA cohorts, we obtained 97.0% and 93.3% sample-wise genotyping call rates (Figure 2), respectively.
4. *High detection rate*. The detection phase of ensembleCNV utilizes multiple underlying callers, and ensures a high detection rate (i.e., ability to detect and genotype more CNVRs). For example, in the FA cohort, ensembleCNV called 19,695 CNVRs, more than the 10,200 CNVRs called by iPattern; in the meantime, ensembleCNV achieved a higher rate of consistency for duplicate pairs (98.6% vs. 90.5%; Figure 2) and fewer Mendelian errors (13 vs. 45; Figure 3).
5. *Improved calling quality and functional interpretability*. The boundary refinement of ensembleCNV both improves the CNV calling quality and downstream functional interpretability.
6. *Prepared for CNV-GWAS*. EnsembleCNV outputs aligned CNVRs and CNV genotype matrix with similar format and properties as SNP genotype matrix (e.g. PLINK format(33)), ready for association testing.
7. *Identification and elimination of batch effects*. Proper ways to handle batch effects are prerequisite to produce high-quality CNV signals. However, this is often overlooked and rarely addressed in a proper way by existing methods. We proposed steps to address this issue at raw data level from the beginning of the pipeline. In the FA dataset, we identified and eliminated batch effects to a considerable extent (Supplementary Results and Supplementary Figures S1-S3 and S11).

CNVs in a given individual can be inherited and *de novo*. We found, when accurately typed, CNVs are overwhelmingly inherited (98.1%), following simple Mendelian inheritance, which is consistent with previous reports.(7) Importantly, we found that inherited CNVs are often not well tagged by nearby SNPs. In the STARNET and FA studies, only 4.4% and 7.3% of CNVRs were tagged by nearby SNPs at r^2^ ≥ 0.6 (Figure 6), emphasizing the need for CNV-GWAS to capture at least part of the missing heritability of human diseases. Further, CNVs are of great functional relevance. Among the unique variants documented in the NHGRI-EBI GWAS catalog, 23.97% (10,530 variants) were inside CNVRs detected in the STARNET and/or FA studies, and 485 variants were inside CNVRs with null (i.e., CN = 0) genotype.

An enormous amount of raw SNP array data generated from large GWASs has not been comprehensively investigated in terms of CNVs, leaving a large gap of CNV genotyping and CNV-based association analyses. To address this gap, we systematically benchmarked the performance of ensembleCNV on large SNP array datasets, suggesting its applicability for CNV-GWAS. In this context, it should be noted that the ensembleCNV framework can be extended to next-generation sequencing (NGS) data (34), where multiple NGS-based CNV calling tools have been proposed (e.g., GenomeSTRiP (35,36) and LUMPY (37)) and their individual call sets can be aggregated in a similar way. Extension of ensembleCNV to NGS data would address the limitations of CNV detection using SNP array data. For example, using the boundary refinement functionality of ensembleCNV, but limited by the pre-specified distribution of probes in a SNP array, the boundaries of CNVs can only be approximated as being somewhere between neighboring probes. By aggregating signals around CNV boundaries (i.e., break point) from split reads and discordant read pairs across multiple individuals, the boundary location may be increased to base-pair resolution (34). Using NGS data with sufficient coverage, we would also alleviate the signal saturation limitation in SNP array data, where we cannot reliably distinguish CN = 3 vs. CN ≥ 4 genotypes at many CNVRs (35,36).

In the 1000 Genomes Project data, some CNVs are more frequent (i.e., frequency ≥ 1%); which are also termed copy number polymorphisms or CNPs (38-40). In the FA data, indicating a high sensitivity for CNPs, we were able to detect and genotype 85% of these frequent CNVs using ensembleCNV. Indeed, the ability to accurately detect and genotype CNPs is particularly important from the perspective of successfully performing CNV-GWAS. The relative contribution of rare and common variants to genetic variation can be measured as a fraction of the number of loci that differ in copy-number between any two unrelated individuals. In a recent analysis, > 90% of the loci were observed to differ in copy-number between pairs of individuals among involved CNPs, and ~80% involved common CNPs (with minor allele frequency >5%) (38-40). This indicates that a large fraction of the copy-number differences between any two individuals arise from a limited set of common polymorphisms (38-40), analogous to an earlier observation that the largest component of human sequence variation (at fine scale) arises from common SNPs. In the FA cohort, 15.8% of the CNVs detected belong to CNPs in which the same mutant allele exists in multiple pedigrees. It should be noted that the detection rate of CNVs is influenced by the density of SNP array probes. Our working definition requires at least five probes per CNV, and SNP arrays with higher probe density as well as specifically designed CNV probes, or next generation sequencing (NGS), may potentially detect even smaller CNVs.

In conclusion, we proposed and benchmarked ensembleCNV, a novel method for CNV calling and genotyping. It makes highly reproducible and accurate CNV calls, obtains a high call rate, and pinpoints CNV boundaries reliably. Importantly, this high performance is not achievable by simply taking the intersection or union of the call sets from individual callers. Our tool is freely available at https://github.com/HaoKeLab/ensembleCNV. Given the amount of SNP array data that has been generated in large GWASs of many diseases and traits, we believe ensembleCNV is a powerful and timely tool to quantify CNV s on existing data, and to investigate the contribution of CNVs to human disease predisposition.

## AVAILABILITY

The source code for ensembleCNV is freely available on GitHub at https://github.com/HaoKeLab/ensembleCNV. The food allergy dataset is available upon reasonable request to the principal investigator, Dr. Xiaobin Wang. The STARNET dataset is available at dbGaP under study accession number: phs001203.v1.p1. The 1000 Genomes Project structural variant data is available at ftp://ftp-trace.ncbi.nih.gov/1000genomes/ftp/phase3/integratedsvmap/

## SUPPLEMENTARY DATA

Supplementary Data including supplementary results, methods, 14 figures and 3 tables are available at NAR Online.

## ACKNOWLEDGMENTS

We thank Dr. Peter Chines, who along with Moebius Syndrome Research Consortium, gave advice.

## FUNDING

This work is partially supported by National Institutes of Health [1R41DA042464-01, 1U01HD079068]; National Natural Science Foundation of China [Grant No. 21477087, 91643201]; the Ministry of Science and Technology of China [Grant No. 2016YFC0206507]; and the Transatlantic Networks of Excellence Award from Foundation Leducq [12CVD02].

## CONFLICT OF INTEREST

The authors declare no conflict of interest.

## References

1. Eichler, E.E., Flint, J., Gibson, G., Kong, A., Leal, S.M., Moore, J.H. and Nadeau, J.H. (2010) Missing heritability and strategies for finding the underlying causes of complex disease. Nature reviews. Genetics, 11, 446–450.

2. Sebat, J., Lakshmi, B., Troge, J., Alexander, J., Young, J., Lundin, P., Maner, S., Massa, H., Walker, M., Chi, M. et al. (2004) Large-scale copy number polymorphism in the human genome. Science, 305, 525–528.

3. Henrichsen, C.N., Chaignat, E. and Reymond, A. (2009) Copy number variants, diseases and gene expression. Hum Mol Genet, 18, R1–8.

4. Chiang, C., Scott, A.J., Davis, J.R., Tsang, E.K., Li, X., Kim, Y., Hadzic, T., Damani, F.N., Ganel, L., Consortium, G.T. et al. (2017) The impact of structural variation on human gene expression. Nat Genet, 49, 692–699.

5. Mefford, H.C. and Eichler, E.E. (2009) Duplication hotspots, rare genomic disorders, and common disease. Curr Opin Genet Dev, 19, 196–204.

6. Cooper, G.M., Zerr, T., Kidd, J.M., Eichler, E.E. and Nickerson, D.A. (2008) Systematic assessment of copy number variant detection via genome-wide SNP genotyping. Nat Genet, 40, 1199–1203.

7. McCarroll, S.A., Kuruvilla, F.G., Korn, J.M., Cawley, S., Nemesh, J., Wysoker, A., Shapero, M.H., de Bakker, P.I., Maller, J.B., Kirby, A. et al. (2008) Integrated detection and population-genetic analysis of SNPs and copy number variation. Nat Genet, 40, 1166–1174.

8. Pinto, D., Darvishi, K., Shi, X., Rajan, D., Rigler, D., Fitzgerald, T., Lionel, A.C., Thiruvahindrapuram, B., Macdonald, J.R., Mills, R. et al. (2011) Comprehensive assessment of array-based platforms and calling algorithms for detection of copy number variants. Nat Biotechnol, 29, 512–520.

9. Wang, K., Li, M., Hadley, D., Liu, R., Glessner, J., Grant, S.F., Hakonarson, H. and Bucan, M. (2007) PennCNV: an integrated hidden Markov model designed for high-resolution copy number variation detection in whole-genome SNP genotyping data. Genome Res, 17, 1665–1674.

10. Colella, S., Yau, C., Taylor, J.M., Mirza, G., Butler, H., Clouston, P., Bassett, A.S., Seller, A., Holmes, C.C. and Ragoussis, J. (2007) QuantiSNP: an Objective Bayes Hidden-Markov Model to detect and accurately map copy number variation using SNP genotyping data. Nucleic Acids Res, 35, 2013–2025.

11. Olshen, A.B., Venkatraman, E.S., Lucito, R. and Wigler, M. (2004) Circular binary segmentation for the analysis of array-based DNA copy number data. Biostatistics, 5, 557–572.

12. Tibshirani, R. and Wang, P. (2008) Spatial smoothing and hot spot detection for CGH data using the fused lasso. Biostatistics, 9, 18–29.

13. Zhang, Z., Lange, K., Ophoff, R. and Sabatti, C. (2010) Reconstructing DNA Copy Number by Penalized Estimation and Imputation. Ann Appl Stat, 4, 1749–1773.

14. Wang, H., Veldink, J.H., Blauw, H., van den Berg, L.H., Ophoff, R.A. and Sabatti, C. (2009) Markov Models for inferring copy number variations from genotype data on Illumina platforms. Hum Hered, 68, 1–22.

15. Diskin, S.J., Li, M., Hou, C., Yang, S., Glessner, J., Hakonarson, H., Bucan, M., Maris, J.M. and Wang, K. (2008) Adjustment of genomic waves in signal intensities from whole-genome SNP genotyping platforms. Nucleic Acids Res, 36, e126.

16. Zhang, Z., Lange, K. and Sabatti, C. (2012) Reconstructing DNA copy number by joint segmentation of multiple sequences. BMC Bioinformatics, 13, 205.

17. Zhang, N.R., Siegmund, D.O., Ji, H. and Li, J.Z. (2010) Detecting simultaneous changepoints in multiple sequences. Biometrika, 97, 631–645.

18. Hong, X., Hao, K., Ladd-Acosta, C., Hansen, K.D., Tsai, H.J., Liu, X., Xu, X., Thornton, T.A., Caruso, D., Keet, C.A. et al. (2015) Genome-wide association study identifies peanut allergy-specific loci and evidence of epigenetic mediation in US children. Nat Commun, 6, 6304.

19. Franzen, O., Ermel, R., Cohain, A., Akers, N.K., Di Narzo, A., Talukdar, H.A., Foroughi-Asl, H., Giambartolomei, C., Fullard, J.F., Sukhavasi, K. et al. (2016) Cardiometabolic risk loci share downstream cis- and trans-gene regulation across tissues and diseases. Science, 353, 827–830.

20. Sudmant, P.H., Rausch, T., Gardner, E.J., Handsaker, R.E., Abyzov, A., Huddleston, J., Zhang, Y., Ye, K., Jun, G., Fritz, M.H. et al. (2015) An integrated map of structural variation in 2,504 human genomes. Nature, 526, 75–81.

21. Guo, Y., He, J., Zhao, S., Wu, H., Zhong, X., Sheng, Q., Samuels, DC., Shyr, Y. and Long, J. (2014) Illumina human exome genotyping array clustering and quality control. NatProtoc, 9, 2643–2662.

22. Dempster, A.P., Laird, N.M. and Rubin, D.B. (1977) Maximum likelihood from incomplete data via the EM algorithm Journal of the Royal Statistical Society. Series B (Methodological), 39, 1–38.

23. DePristo, M.A., Banks, E., Poplin, R., Garimella, K.V., Maguire, J.R., Hartl, C., Philippakis, A.A., del Angel, G., Rivas, M.A., Hanna, M. et al. (2011) A framework for variation discovery and genotyping using next-generation DNA sequencing data. Nat Genet, 43, 491–498.

24. Korn, J.M., Kuruvilla, F.G., McCarroll, S.A., Wysoker, A., Nemesh, J., Cawley, S., Hubbell, E., Veitch, J., Collins, P.J., Darvishi, K. et al. (2008) Integrated genotype calling and association analysis of SNPs, common copy number polymorphisms and rare CNVs. Nat Genet, 40, 1253–1260.

25. Zhang, Q., Ding, L., Larson, D.E., Koboldt, D.C., McLellan, M.D., Chen, K., Shi, X., Kraja, A., Mardis, E.R., Wilson, R.K. et al. (2010) CMDS: a population-based method for identifying recurrent DNA copy number aberrations in cancer from high-resolution data. Bioinformatics, 26, 464–469.

26. Fromer, M., Moran, J.L., Chambert, K., Banks, E., Bergen, S.E., Ruderfer, D.M., Handsaker, R.E., McCarroll, S.A., O’Donovan, M.C., Owen, M.J. et al. (2012) Discovery and statistical genotyping of copy-number variation from whole-exome sequencing depth. Am J Hum Genet, 91, 597–607.

27. Genomes Project, C., Auton, A., Brooks, L.D., Durbin, R.M., Garrison, E.P., Kang, H.M., Korbel, J.O., Marchini, J.L., McCarthy, S., McVean, G.A. et al. (2015) A global reference for human genetic variation. Nature, 526, 68–74.

28. Welter, D., MacArthur, J., Morales, J., Burdett, T., Hall, P., Junkins, H., Klemm, A., Flicek, P., Manolio, T., Hindorff, L. et al. (2014) The NHGRI GWAS Catalog, a curated resource of SNP-trait associations. Nucleic Acids Res, 42, D1001–1006.

29. Solovieff, N., Cotsapas, C., Lee, P.H., Purcell, S.M. and Smoller, J.W. (2013) Pleiotropy in complex traits: challenges and strategies. Nat Rev Genet, 14, 483–495.

30. Gratten, J. and Visscher, P.M. (2016) Genetic pleiotropy in complex traits and diseases: implications for genomic medicine. Genome Med, 8, 78.

31. Zhang, J., Peng, S., Cheng, H., Nomura, Y., Di Narzo, A.F. and Hao, K. (2017) Genetic Pleiotropy between Nicotine Dependence and Respiratory Outcomes. Sci Rep, 7, 16907.

32. Wang, Z., McGlynn, K.A., Rajpert-De Meyts, E., Bishop, D.T., Chung, C.C., Dalgaard, M.D., Greene, M.H., Gupta, R., Grotmol, T., Haugen, T.B. et al. (2017) Meta-analysis of five genome-wide association studies identifies multiple new loci associated with testicular germ cell tumor. Nat Genet, 49, 1141–1147.

33. Purcell, S., Neale, B., Todd-Brown, K., Thomas, L., Ferreira, M.A., Bender, D., Maller, J., Sklar, P., de Bakker, P.I., Daly, M.J. et al. (2007) PLINK: a tool set for whole-genome association and population-based linkage analyses. Am J Hum Genet, 81, 559–575.

34. Medvedev, P., Stanciu, M. and Brudno, M. (2009) Computational methods for discovering structural variation with next-generation sequencing. Nat Methods, 6, S13–20.

35. Handsaker, R.E., Korn, J.M., Nemesh, J. and McCarroll, S.A. (2011) Discovery and genotyping of genome structural polymorphism by sequencing on a population scale. Nat Genet, 43, 269–276.

36. Handsaker, R.E., Van Doren, V., Berman, J.R., Genovese, G., Kashin, S., Boettger, L.M. and McCarroll, S.A. (2015) Large multiallelic copy number variations in humans. Nat Genet, 47, 296–303.

37. Layer, R.M., Chiang, C., Quinlan, A.R. and Hall, I.M. (2014) LUMPY: a probabilistic framework for structural variant discovery. Genome Biol, 15, R84.

38. McCarroll, S.A. and Altshuler, D.M. (2007) Copy-number variation and association studies of human disease. Nat Genet, 39, S37–42.

39. McCarroll, S.A. (2008) Extending genome-wide association studies to copy-number variation. Hum Mol Genet, 17, R135–142.

40. Girirajan, S., Campbell, C.D. and Eichler, E.E. (2011) Human copy number variation and complex genetic disease. Annu Rev Genet, 45, 203–226.

